# α-Synuclein Seeding in Dopaminergic Neurons Transplanted into the Putamen of Parkinson’s Disease Patients

**DOI:** 10.64898/2026.09.22.751740

**Authors:** Jayda B. Duvernay, Bryan A. Killinger, Solji G. Choi, Tyler Tittle, Atousa Bahrami, Marla E. Tharp, Gabriela Mercado, Yaping Chu, Patrik Brundin, Jeffrey H. Kordower

## Abstract

Healthy embryonic neurons grafted into the putamen of Parkinson’s disease (PD) patients start to develop Lewy pathology 10 years after transplantation. It remains unclear whether this pathology is initiated by the spread of host-derived α-synuclein (αsyn) aggregates which seed aggregation in the grafted cells or arises independently from a hostile PD environment (e.g., chronic inflammation).

To distinguish these possibilities, we identified αsyn seeding sites in the human brain using a multiplex *in situ* seed immunodetection (isSID) assay. In PD and dementia with Lewy bodies (DLB), isSID labeled a subset of Lewy pathology (LP) as puncta along neurites and within inclusions. In multiple system atrophy (MSA), isSID labeled Papp-Lantos bodies and microglia. In progressive supranuclear palsy (PSP), isSID labeled tufted astrocytes. isSID labeled neuromelanin in the substantia nigra independent of diagnosis. In 16- and 27-year intraputamenal neuronal grafts, isSID labeled within and proximal to Lewy pathology, as well as outside the grafted tissue. Younger grafts (18 months and 4 years) lacked Lewy pathology. In the 18-month graft, isSID labeled grafted neurons; at 4 years, isSID was confined to amorphous deposits in the graft and was absent from tyrosine hydroxylase-positive grafted neurons.

In conclusion, αsyn seeding is detectable in grafted neurons before and during LP formation. Because seeding accompanies melanization, but is not explained by melanization alone, these findings support host-derived seeds as a contributor to graft pathology.

## Introduction

Immature dopamine neurons grafted into the putamen innervate the surrounding host tissue and survive many years following implantation [42, 44, 48, 55]. Over several decades, brains from patients who have undergone transplantation surgery have been examined microscopically after the patients have died. In the published cases, the graft survival times have varied greatly, ranging from 18 months to 24 years. In young grafts after implantation, diffuse cytosolic αsyn accumulation is observed, and as grafts age (>10 years), a subset of neurons between 5%-30% developed Lewy pathology (LP) that is indistinguishable from the pathology observed in the host neurons[42, 45, 48, 49]. The LP in grafted neurons is filamentous, ubiquitinated and thioflavin positive, and takes on a variety of appearances from compact ring-like to loose meshwork[49]. Infiltrating microglia and reactive glia[46] have been observed before obvious αsyn accumulation or development of LP in grafts, suggesting the possibility that an inflammatory pathogenic microenvironment triggered αsyn accumulation/misfolding and subsequent LP formation[56]. However, similar grafts in Huntington’s patients also show inflammatory responses, but do not appear to develop intracellular LP[24, 39]. Extracellular Huntington’s aggregates can be seen within these grafts, but this may be due to degeneration of cortical cells innervating the graft with the aggregates leaching from degenerating neurites [15, 16]. Together, these findings suggest that a pathological inflammatory environment alone is insufficient to induce LP[5]. Indeed, inflammatory processes in fetal dopamine grafts occur quickly, but LP takes at least a decade post-grafting to occur[56]. Rather, LP in grafted neurons appears to result from the spread of αsyn from the host brain to the grafts. However, the exact mechanism underlying LP formation in the grafted neurons remains unclear.

The αsyn spreading mechanism has been investigated in cell culture models and in experimental animals[51, 54, 63], but direct evidence of spreading in the human brain remains sparse. Braak hypothesized that αsyn spreads as PD progresses based on the observation that, at least for PD, αsyn pathology progresses in a stereotypical caudal-rostral pattern with brainstem and olfactory structures being affected early and severely[2–4, 7, 37]. Observations that healthy neurons grafted into the PD putamen developed αsyn pathology provided evidence that αsyn pathology spread can occur in the human PD brain[5, 42, 48]. Some preclinical studies support the spread mechanism; injections of brain material or synthetic αsyn filaments (αsyn preformed fibrils, “PFFs”) into the rodent and nonhuman primate brain result in αsyn inclusion formation at the site of injection and amongst first-order neurons[14, 35, 62, 66], and other factors, such as transgene overexpression, promote seeding and spread[1, 51, 58]. Moreover, mice that are null mutants for the *Snca* do not exhibit any αsyn pathology after injection of αsyn PFFs[51, 65]. While PFFs have been a useful model to study αsyn aggregation and spread, there are still questions concerning whether spread occurs and is a driving factor in human disease[40]. Although seeds of misfolded αsyn are likely to be involved in spreading, the origins of seeds in human brain and the mechanisms behind de novo seed formation are unclear. For spread to occur, αsyn beta-sheet conformations are required for templated αsyn aggregation, with primary nucleation at filament ends or secondary nucleation at the aggregate surface driving the aggregation process[6, 26, 34, 67]. Although pathological αsyn aggregates contain beta-sheet αsyn conformations[18, 23, 29, 47, 53, 61], steric constraints of inclusions that restrict access to a nucleating surface, likely limit where seeding occurs in diseased neurons. It is unclear where the seeding occurs in cells that develop αsyn pathology.

Several methods have been developed for the detection of αsyn seeding activity in biological samples. The seed amplification assay (SAA) is an exquisitely sensitive method for detecting seeds in biological fluids[19, 59, 60], but it does not reveal the cellular location and origins of seeding. Recently, a variant of SAA was adapted for use *in situ*, allowing for the detection of αsyn seeding and determination of seeding location in human brain formalin-fixed specimens[57]. One of these techniques, *in situ* seeding immunodetection (isSID), demonstrated active αsyn seeding in neurons and glia in various synucleinopathies[57]. isSID revealed potential cross-seeding of αsyn with tau tangles and amyloid plaques, supporting the role of co-pathologies’ influence on αsyn aggregate formation[57]. Notably, isSID has the potential to identify active points of seeding in tissues, as opposed to traditional neuropathological markers like phosphorylated αsyn (pSer129), which are a proxy of aggregates in the post-mortem human brain[11, 25]. In PFF seeding models, isSID reveals discontinuous active seeding in pathology bearing cells[10], suggesting active seeding and αsyn inclusions are spatially separated. Determining the location of seeding within cells/tissue is critical for understanding mechanisms contributing to the development of synucleinopathy.

Here, to determine the precise location of αsyn seeding in the human brain, we adapted isSID with our tyramide multiplex labeling approach (https://dx.doi.org/10.17504/protocols.io.261gey7bov47/v1), allowing for enhanced detection and applied it to the human synucleinopathy brain. Then we used this technique to determine whether active seeding occurs in originally healthy neurons implanted into the PD putamen.

## Materials and methods

### Tissues

Rodent tissues were obtained from wild-type C57BL/6 and αsyn knockout (C57BL/6N-SncatmMjff/J, Jackson Labs) mice maintained under institutionally approved IACUC protocols. Animals were anesthetized with ketamine/xylazine and transcardially perfused with PBS (pH 7.4), followed by 4% paraformaldehyde (PFA) in PBS. Brains were collected and post-fixed overnight at 4°C in 4% PFA. Tissues were then cryoprotected in a graded sucrose series (10%, 20%, and 30% w/v in PBS) and sectioned coronally at 40 µm using a freezing-stage sliding knife microtome. Postmortem human brain tissue was obtained from the Rush Movement Disorders Brain Bank and the Arizona State University–Banner Neurodegenerative Disease Research Center with approval of the respective institutional review boards. The study was performed in accordance with the 1964 Declaration of Helsinki and its later amendments. Informed consent for brain donation and research use was obtained from all donors. Cases included those with a primary clinical diagnosis of multiple system atrophy (MSA), dementia with Lewy bodies (DLB), and Parkinson’s disease (PD) (see Table 1 for case characteristics). Graft cases of varied age were used including 18 months, 4 years, 16 years, and 27 years after implantation, and, except for the 27-year case, their characterization was previously described[13, 44, 45]. All tissue handling procedures were carried out according to a previously established protocol[12]. In brief, coronal slabs (∼2 cm thick) were immersion-fixed in 4% PFA prepared in 0.1 M phosphate-buffered saline (PBS; pH 7.4) at 4°C for seven days. Following fixation, slabs were gradually equilibrated in a cryoprotectant solution consisting of PBS supplemented with 2% dimethyl sulfoxide and 20% glycerol. Regions of interest were then dissected and sectioned at 40 µm using a freezing-stage sliding knife microtome (American Optical). Sections were stored in cryoprotectant (30% sucrose, 30% ethylene glycol in PBS) at −20°C until further use.

**Table 1.** Characteristics of synucleinopathy and non-synucleinopathy patients. NA = data not available.

| Sample # | Sex | Age at death (years) | Hoehn and Yahr scale | Disease duration (years) | Clinical diagnosis | Pathological diagnosis |
| --- | --- | --- | --- | --- | --- | --- |
| 1 | F | 75 | 4 | 4 | MSA-P | MSA-P |
| 2 | F | 65 | 5 | 10 | MSA-P | NA |
| 3 | M | 55 | 4 | 6 | MSA-P | NA |
| 4 | M | 67 | 5 | 8 | MSA | NA |
| 5 | M | 74 | NA | 9 | PD | Corticobasal degeneration with parkinsonism |
| 6 | NA | NA | NA | NA | PD | NA |
| 7 | F | 87 | 4 | 3 | Dementia; PD secondary | NA |
| 8 | M | 69 | NA | 4 | PD | NA |
| 9 | NA | NA | NA | NA | PD | LBD |
| 10 | M | 76 | NA | 11 | PD | NA |
| 11 | M | 70 | 4 | 5 | PD/PSP | NA |
| 12 | NA | NA | NA | NA | NS | Corticobasal degeneration; Alzheimer's |
| 13 | NA | NA | NA | NA | NS | Alzheimer's; FXTAS-AD |
| 14 | NA | NA | NA | NA | NS | PSP; FXTAS-AD |

**Table 2.**
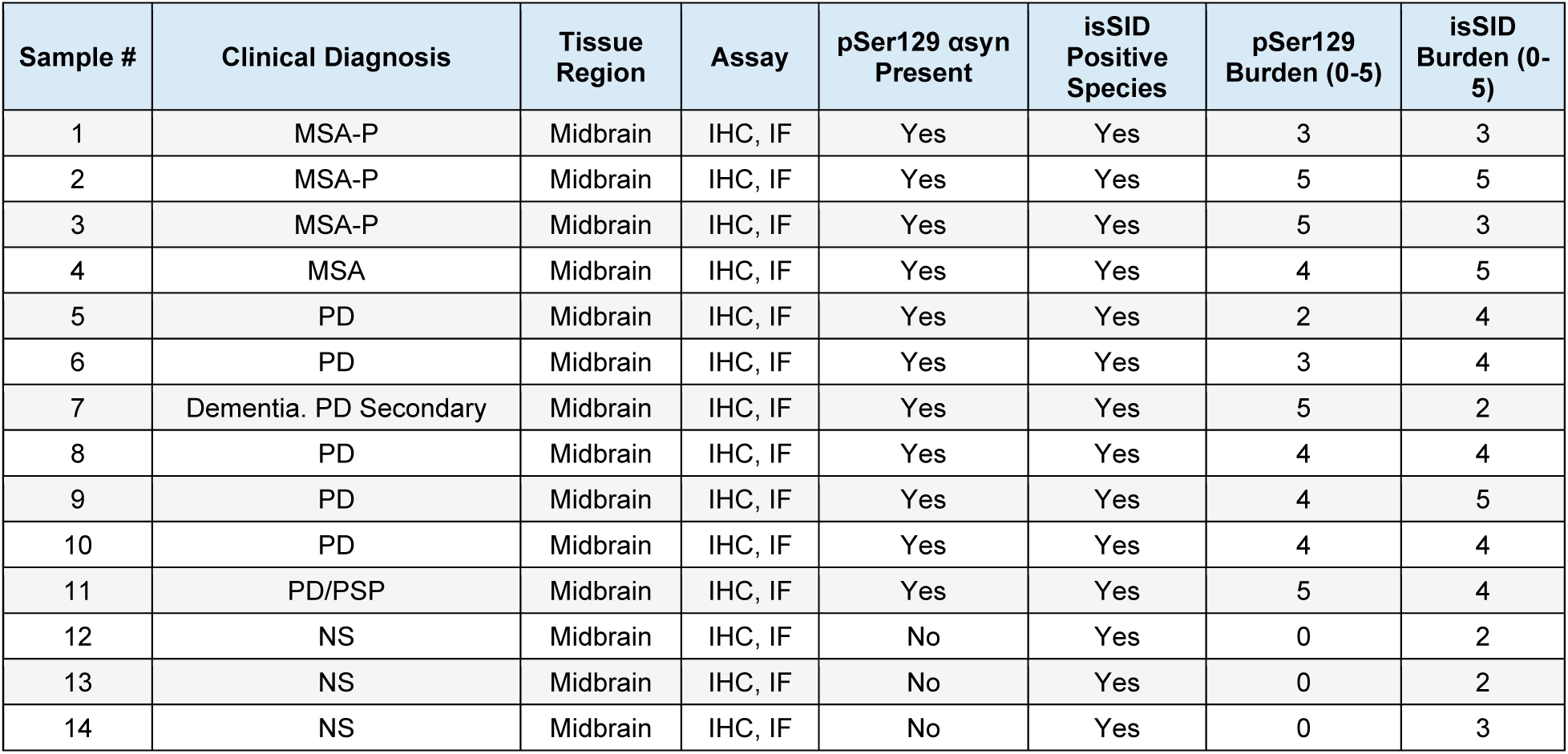
Summary of results across synucleinopathy and non-synucleinopathy (NS) cohort.

**Table 3.** Summary of results across putamen grafts cohort.

| Sample # | Tissue Region | Assay | Host Seeding | isSID in grafted TH+ neurons | pSer129 in TH+ neurons |
| --- | --- | --- | --- | --- | --- |
| 18-month graft | Striatum | IF | Yes | Yes | No |
| 4-year graft | Striatum | IF | Yes | No | No |
| 16-year graft | Striatum | IF | Yes | Yes | Yes |
| 27-year graft | Striatum | IF | Yes | Yes | Yes |

### In situ seed immunodetection (isSID)

We adapted a previously described protocol[10, 57]. Free-floating brain sections were washed three times in dilution media (DM; 50 mM Tris–HCl, pH 7.4, 150 mM NaCl, 0.05% Triton X–100) and twice in Tris-buffered saline (TBS; pH 7.6). Endogenous peroxidase activity was quenched by incubating sections in 0.3% hydrogen peroxide prepared in TBS for 30 min. Heat-induced antigen retrieval (HIAR) was performed in citrate buffer (10 mM citric acid, 0.05% Tween–20, pH 6.0) at 90°C for 10 min, followed by immediate cooling on ice until the solution reached room temperature. Sections were then washed three times in phosphate-buffered saline (PBS; pH 7.4) and equilibrated in preconditioning buffer containing 100 mM PIPES for 1 h. Sections were then incubated overnight at 37°C on a thermomixer with either human His-αsyn (Acro Biosystems, Cat# ALNH52H8) or mouse His-αsyn (Acro Biosystems, Cat# ALN-M52H6) at a final concentration of 0.025 mg/ml in 100 mM PIPES buffer. For His-GFP controls, recombinant His-GFP (ThermoFisher Scientific, Cat# A42613) was reconstituted in sterile water to 0.2 mg/mL and subsequently diluted in PIPES buffer to match the final molar concentration of His-αsyn. Following incubation, sections were washed three times in TBS and fixed in 4% PFA for 10 min. After fixation, sections were washed in PBST (0.5% Tween–20 in PBS, pH 7.4). Non-specific binding was blocked by incubating sections for 1 h in blocking buffer consisting of TBS supplemented with 0.5% Triton X–100 and 10% normal serum. Sections were then incubated overnight at 4°C with mouse anti-His primary antibody (ThermoFisher Scientific Cat# MA1-21315-HRP, RRID: AB_2536989) diluted 1:5,000 or rabbit anti-His primary antibody (Proteintech Cat# 1000-0-AP, RRID: AB_11042316) diluted 1:2,700 in blocking buffer. The following day, sections were washed three times in PBST and incubated with biotinylated horse anti-mouse secondary antibody (Vector Laboratories Cat# BA-2000, RRID: AB_2313581) or goat-anti rabbit (Vector Laboratories Cat# BA-1000, RRID: AB_2313606) diluted 1:200 in blocking buffer for 1 h. Sections were washed again in PBST and incubated with avidin–biotin complex (ABC) reagent (Vector Laboratories Cat# PK-6100, RRID: AB_2336819) according to the manufacturer’s instructions. Tissues were subsequently washed three times in PBST and once in sodium acetate buffer (0.2 M imidazole, 1.0 M sodium acetate buffer, pH 7.2). Nickel-enhanced DAB visualization was then performed according to a previously established protocol (dx.doi.org/10.17504/protocols.io.rm7vzx725gx1/v1). Sections were then rinsed with sodium acetate buffer followed by PBS and mounted onto charged Superfrost Plus Microscope Slides (Fisher Scientific, Cat# 22-034-979) and then dried completely. Purified methyl green (Sigma-Aldrich, Cat# 67060) was used as a counterstain. Sections were dehydrated, cleared in xylenes, and coverslipped using Cytosol XYL mounting medium (Epredia, Ref# 8312-4)

### TSA multiplex Immunofluorescence (IF)

isSID was performed as previously described above. Following incubation with the ABC reagent and subsequent washes in PBST, tissue sections were washed once in borate buffer (0.05 M borate buffer, pH 8.5) and incubated for 30 min in freshly prepared fluorescent tyramide (FT) working solution (borate buffer, 0.003% hydrogen peroxide, 5 µM CF568 (Biotium, Cat# 92173), protected from light. Sections were then stored overnight in PBS. The following day, sections were washed twice in DM, and residual peroxidase activity was quenched by incubating sections for 1 h in blocking buffer (3% normal serum and 0.4% Triton X-100 in DM) supplemented with 0.3% hydrogen peroxide and 0.1% sodium azide. Sections were subsequently incubated overnight at 4°C with pSer129 αsyn antibody (Abcam “EP1536Y” Cat# ab51253, RRID: AB_869973) diluted 1:50,000, pS396 Tau antibody (Abcam “EPR2731” Cat# ab109390, RRID: AB_10860822) diluted 1:500, TH antibody (Abcam Cat# ab76442, RRID: AB_1524535), or IBA1 antibody (FUJIFILM Wako Pure Chemical Corporation Cat# 019-19741, RRID: AB_839504) diluted 1:1000 in blocking buffer. For sections in which the ABC reagent was used twice, an avidin–biotin blocking kit (Vector Laboratories Cat# SP-2001, RRID: AB_2336231) was applied according to manufacturer’s instructions prior to primary antibody incubation. The next day, sections were washed three times in DM and incubated with a goat anti-rabbit HRP-conjugated secondary antibody (ThermoFisher Scientific Cat# 31462, RRID: AB_228338) diluted 1:1,000 or biotinylated goat anti-chicken secondary antibody (Vector Laboratories Cat# BA-9010, RRID: AB_2336114) diluted 1:200 in blocking buffer for 1h. Sections were then washed three times in DM and once in borate buffer before incubation for 30 min in freshly prepared FT working solution (borate buffer, 0.003% hydrogen peroxide, 5 µM CF488 (Biotium, Cat# 92171), protected from light. For triple labeling, the blocking, primary antibody, and secondary antibody steps were repeated as appropriate for the third target. Sections were subsequently washed in DM and once in borate buffer before incubation for 30 min in freshly prepared FT working solution (borate buffer, 0.003% hydrogen peroxide, 5 µM CF640R (Biotium, Cat# 92175), protected from light. Sections were washed twice in PBS and incubated for 20 min with DAPI (MilliporeSigma, 26829810MG, Cat# 26-829-810MG, for stock, reconstituted in ddH2O at a concentration 5mg/mL) diluted 1:2,000 in Milli-Q water. After a brief rinse in PBS, sections were mounted on glass slides, dried for 25 min, and covered with #1.5 glass coverslips using FluoroShield mounting medium (MilliporeSigma, F6182-20ML, Cat# 01-258-928).

### Immunohistochemistry (IHC)

Free-floating brain sections were washed three times in DM, followed by heat-induced antigen retrieval (HIAR) in sodium citrate buffer (10 mM sodium citrate, 0.05% Tween–20, pH 6.0) at 90°C for 30 min and then immediately cooled on ice until the solution reached room temperature. Tissue sections were then washed twice in DM. Endogenous peroxidase activity was quenched by incubating sections for 1 h in blocking buffer (3% normal serum and 0.4% Triton X–100 in DM) supplemented with 0.3% hydrogen peroxide and 0.1% sodium azide. Sections were incubated overnight at 4°C with pSer129 αsyn antibody diluted 1:50,000 or Phospho-Tau (ThermoFisher Cat# MN1020, RRID: AB_223647) antibody diluted 1:1000 in blocking buffer. The following day, sections were washed three times in DM and incubated with a biotinylated goat anti-rabbit (Vector Laboratories Cat# BA-1000, RRID: AB_2313606) or horse anti-mouse (Vector Laboratories Cat# BA-2000, RRID: AB_2313581) secondary antibody diluted 1:200 in blocking buffer for 1 h. Sections were then washed twice in DM, followed by signal amplification using an avidin–biotin complex (ABC) reagent for 1 h. Tissues were subsequently washed twice in DM and once in sodium acetate buffer. DAB development, counterstaining, clearing, and coverslipping were performed as described above.

### Microscopy and Imaging

All prepared slides were systematically surveyed from end to end using a 10X objective to assess the overall distribution and abundance of signals. Representative regions spanning the observed pathological features were subsequently selected for higher-magnification imaging. Selected regions were imaged using a Nikon A1R confocal microscope controlled with NIS-Elements software (version AR 6.10.03 64 bit). Brightfield images for neuromelanin were acquired on a Zeiss Axio observer 7 with apotome 3 camera and image processed using ZEN 3.10 (ZEN pro).

### Fluorescence microscopy

Large fluorescence overview images were acquired as tiled z-stack scans, followed by denoising and maximum intensity projection (MAX IP) within NIS-Elements. Images used to show three-dimensional volume distribution were acquired as z-stacks and reconstructed using the 3D volume distribution function in NIS-Elements. All remaining fluorescence images were acquired using a 10X, 20X or 60X objective lenses, as either z-stacks followed by 3D deconvolution and then MAX IP or as single-plane images followed by 2D deconvolution. Lookup tables (LUTs) were adjusted for image visualization. To avoid spectral bleed-through fluorescent channels were acquired sequentially. Due to the exceptional fluorescent intensity of tyramide amplification, images were captured at the instrument’s near-minimal operational thresholds (e.g., 0.4% laser power; PMT gain of ∼20).

### Brightfield microscopy

Large brightfield overview images were acquired as single-plane tiled scans using the same microscope platform. Brightfield images acquired using 20X or 60X objective were collected as single-plane images. For figure preparation, images underwent cropping, resizing, and uniform adjustments to brightness and contrast in Adobe Photoshop (version 27.7.0) to improve clarity and presentation. Processed images were subsequently imported into Adobe Illustrator (version 30.4.0) for figure assembly and final layout.

### Slide scoring

Samples were evaluated under a 10X objective for the presence, distribution, and relative abundance of both pSer129- and isSID-positive species. Signal was assessed using a semi-quantitative scoring system ranging from 0 to 5, where 0 indicated the absence of detectable staining or pathology, and increasing scores reflected progressively greater abundance and distribution of labeled species within the evaluated tissue region. A score of 1 was assigned when rare, isolated, or minimally detectable signal was observed; a score of 2 represented low-level signal with limited distribution of labeled species; a score of 3 represented moderate signal burden with more frequent labeling and broader regional involvement; a score of 4 indicated high regional abundance with widespread distribution throughout the evaluated region; and a score of 5 represented extensive signal burden with abundant and widespread labeling throughout the tissue region. Scoring was performed independently for pSer129 and isSID-positive signal.

Initial scoring was performed by an unblinded reviewer using the predefined criteria established above. To assess reproducibility of the scoring approach, slides were randomized and placed in a slide box in a non-sequential order. An independent reviewer was blinded to sample identity, clinical diagnosis, and experimental group during evaluation. Prior to scoring, the reviewer was provided with a representative reference slide illustrating examples spanning the range of observed pSer129 and isSID signal abundance to establish consistent interpretation of the scoring criteria. Following independent scoring by both the primary and blinded reviewer, scores were compared for consistency. Because most assessments demonstrate concordance between reviewers, the scores assigned by the blinded reviewer were used as the final scores. In rare instances where discrepancies occurred between the reviewers, the corresponding samples were re-evaluated, and scores were resolved through discussion and consensus.

## Results

We applied isSID across synucleinopathy, non-synucleinopathy, and grafted brain specimens (See table 1-3 for summary of observations). isSID[10, 57] and similar approaches[52], have raised the prospect of determining the precise location of αsyn seeding in the human brain. Underlying principles of these assays are similar (Fig 1A), his-tagged-recombinant αsyn is incubated with formaldehyde fixed tissues and binds to moderate-affinity sites (1.53 μM αsyn concentration used here) within the tissues (Fig 1A, step 1). Subsequently αsyn binding is detected by concentrating HRP at the site using standard biotinylated secondary antibodies with ABC (Fig 1A. steps 2-4). Colorimetric DAB or fluorescent tyramide protocols permanently labeled the location of αsyn binding for downstream analysis. Early validation (Fig. 1B) demonstrated several critical parameters for isSID, including heat mediated antigen retrieval, which reduced diffuse αsyn binding throughout the grey matter. Secondary detection sensitivity was also critical, with ABC based detection (DAB or tyramide) giving the highest sensitivity. Following optimization, isSID-positive concentric Lewy bodies were apparent in the midbrain (Figure 1B, bottom panels). isSID labeled apparent Papp-Lantos bodies in the MSA brain and LP in the PD/DLB brain, although pSer129 reactivity was more intense, and widespread than isSID (Figure 1C). Comparing synucleinopathy with non-synucleinopathy brain (Fig 1D), we observed strongly isSID reactive Lewy bodies throughout the midbrain, as well as reactivity within pigmented nigral cells (Fig. 1D, Fig. S1). αSyn affinity for neuromelanin [21] and neuromelanin associated lipids[32] has been previously described[68]. Similarly, both mouse and human αsyn bound diffusely to WT and SNCA KO mouse brain (Fig. 1E). This diffuse labeling was substantially reduced when detection was preformed using a secondary antibody from a different host species, suggesting that the initial background was associated, at least in part, with interference from the mouse-derived secondary despite pretreatment with mouse-on-mouse (M.O.M) blocking reagent (Fig. 1E, Fig. S5). Our observations of nonspecific, non-αsyn binding, make isSID-DAB non-ideal for determining αsyn seeding location. Thus, we adopted tyramide labeling strategies allowing us to multiplex isSID with markers of LP (e.g., pSer129) (Fig. 2).

**Figure 1.**
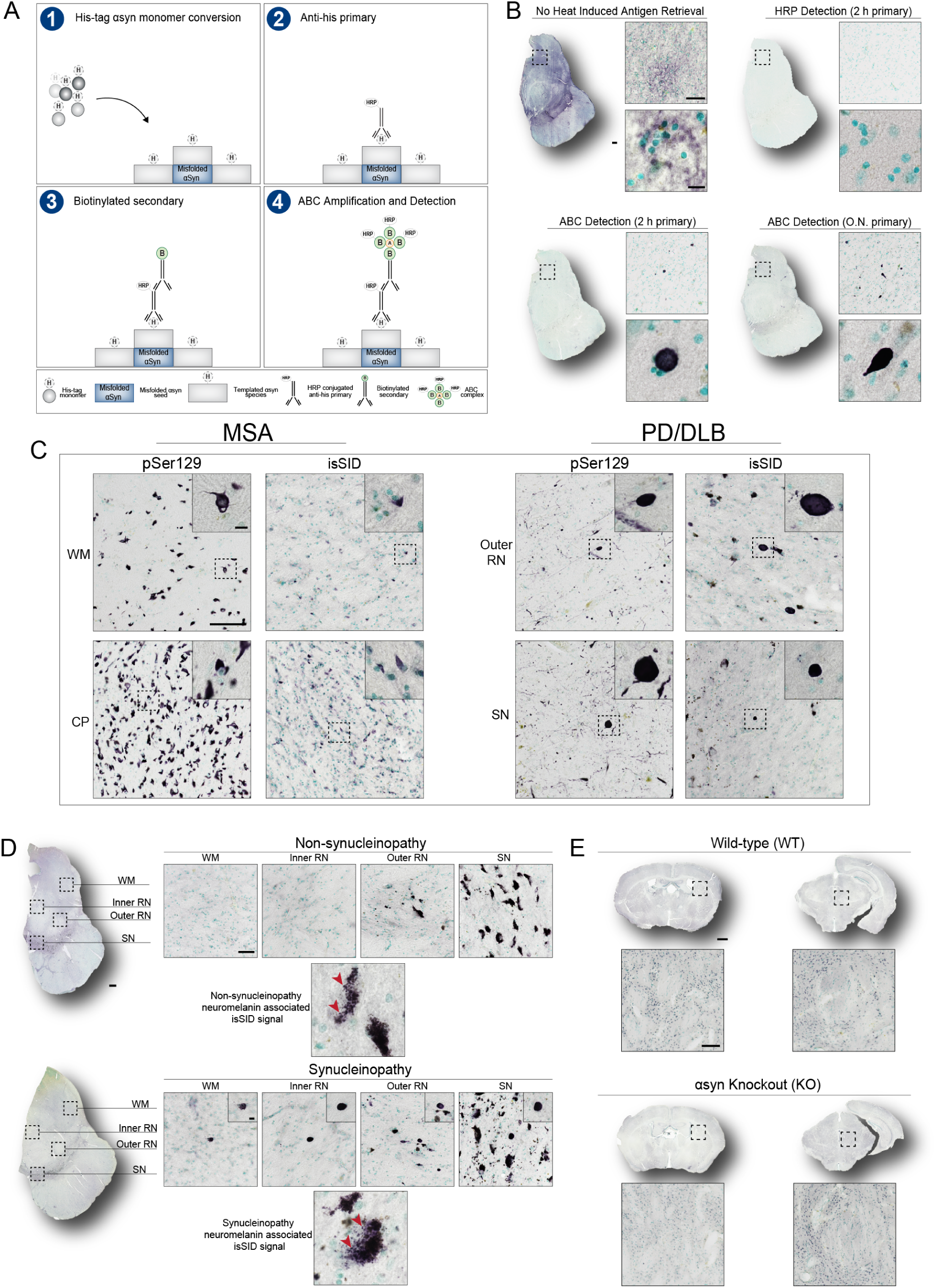
isSID reveals the spatial distribution of seed-competent αsyn in the synucleinopathy brain. **(A)** Schematic illustrating the optimized isSID assay, highlighting major assay steps from substrate incubation through signal amplification. (1) His-tagged αsyn monomers bind to seed-competent surfaces. (2) Then Anti-His binds to His-tagged αsyn assemblies (3) Biotinylated secondary antibody binds to the HRP-conjugated primary antibody. (4) ABC complex binds to biotin and enables signal amplification and detection via DAB or fluorescent labeling. **(B)** Optimization of isSID demonstrating increasing assay sensitivity across tested conditions: (1) no heat-induced antigen retrieval, (2) HRP detection with a 2-hour primary antibody incubation, (3) ABC detection with a 2-hour primary antibody incubation, and (4) ABC detection with an overnight primary antibody incubation which yielded the most sensitive detection method. Scale bars = 10 µm, 100 µm, and 1 mm. **(C)** Representative pSer129 immunostaining and corresponding isSID labeling in the synucleinopathy brain. MSA samples showed more abundant isSID labeling in areas such as white matter (WM) and cerebral peduncle (CP), while PD/DLB samples displayed abundant isSID labeling in areas such as the immediate outer portion of the red nucleus (Outer RN) and substantia nigra (SN) Scale bars = 10 µm and 100 µm. **(D)** Comparison of isSID labeling in non-synucleinopathy and synucleinopathy brain highlighting the absence of Lewy-type isSID signal in the non-synucleinopathy brain. However, the presence of similar punctate isSID signal patterns was observed in the neuromelanin-containing cells regardless of disease diagnosis. Scale bars = 10 µm, 50 µm, and 1 mm. **(E)** Comparison of wild-type and αsyn knockout mice in the striatum and periaqueductal grey. Diffuse grey matter labeling is observed across both conditions and not eliminated by αsyn knockout. Scale bar = 100 µm and 1 mm.

**Figure 2.**
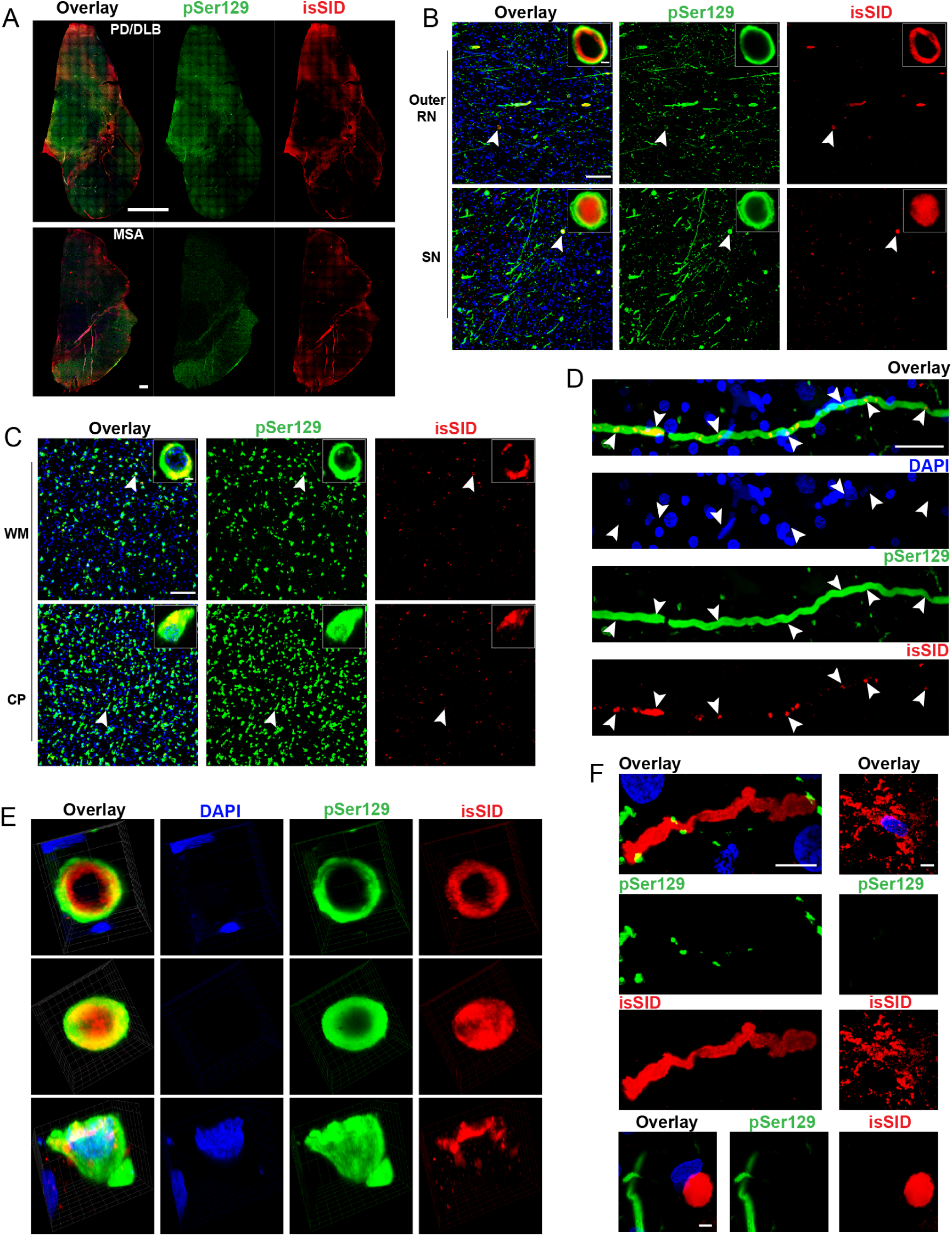
Seed competent αsyn is restricted to a subset of pSer129 pathology. **(A)** 10X overview of PD/DLB and MSA brain sections immunolabeled for pSer129 (green), isSID (red), and DAPI (blue, nuclei). Scale bars = 1 mm and 5 mm. **(B-C)** Low- and high-magnification images from the outer red nucleus (Outer RN) and substantia nigra (SN) in the PD/DLB brain (B) and the white matter (WM) and cerebral peduncle (CP) in the MSA brain (C). isSID signal is detected within Lewy bodies (PD/DLB) and glial cytoplasmic inclusions (MSA). Although isSID-positive seed-competent αsyn represents only a subset of the total pSer129 pathology, its abundance closely mirrors the regional distribution of pSer129 pathology. Scale bars = 5 µm and 100 µm. **(D)** A neuritic process from the PD/DLB brain showing pSer129 immunoreactivity throughout its length, whereas isSID signal is restricted to discrete punctate foci along the axon. Scale bar = 50 µm. **(E)** Three-dimensional volume reconstructions illustrating the distribution of isSID signal within individual αsyn inclusions. Lewy bodies were found to have two main types of seed distribution occurring at either the periphery with reduced labeling in the central core or a more homogenous distribution that filled the entirety of the structure. In contrast, isSID distribution within glial cytoplasmic inclusions were mainly confined to the top portion rather than throughout its entire volume. **(F)** Representative examples of isolated isSID-positive structures lacking detectable pSer129 immunoreactivity, including neurite-like and Lewy body-like structures identified in the PD/DLB brain, and a glial cell-like structure identified in the MSA brain. Scale bars = 5 µm and 10 µm.

Midbrain sections from PD/DLB and MSA brain highlight the overall distribution of pSer129 and isSID signal. In PD/DLB, pSer129 is heavily distributed in the grey matter of the SN, red nucleus and outer reticular nucleus (RN) (Fig. 2A). In MSA, pSer129 is heavy in white matter pontine fibers. isSID overlap with pSer129 was minimal at this magnification, except strong overlap in the SN in PD brain (Fig. 2A, overlap appears yellow). Higher magnification revealed isSID positive inclusions in PD/DLB, and MSA brain (Fig. 2B, C). isSID positive neurites and concentric bodies were seen in the PD/DLB, and glial cytoplasmic inclusions (GCI) in MSA (2D, E). In all specimens tested we observed isSID positive structures that lacked pSer129 (Fig. 2F). Furthermore, we observed pSer129 positive structures that lacked detectable isSID (Fig. 2F). In most instances, isSID-labeling of pSer129-inclusions was discontinuous, either punctate along Lewy neurites (Fig. 2D, arrows), within inner Lewy body rings (Fig. 2E), or discrete punctate within GCIs.

isSID specifically labeled αsyn aggregates (Fig. 2) and showed some affinity for neuromelanin (Fig. 1, Fig. S2), however; depending on the brain specimen, we observed other high affinity αsyn binding without overlapping with αsyn markers (i.e., pSer129). For example, PSP brain that lacked αsyn pathology showed profound isSID labeling in the midbrain (i.e., substantia nigra) (Fig. 3A) that resembled amyloid plaques but were not beta amyloid reactive (Fig. S4). Instead, these large isSID reactive deposits were found in tufted astrocytes, proximal to Tau aggregates, confirmed with multiplex labeling (Fig. 3B) and standard DAB protocols (Fig. 3C). In MSA brain, we observed strong isSID signal in white matter, that did not overlap with pSer129/GCIs. IBA1 staining revealed a close association of isSID and microglia (Figure 3D). Strikingly, in one MSA brain (Sample #4), dense isSID positive microglia nodules were concentrated in cerebral peduncle, consistent with white matter degeneration in MSA-P (Fig. 3D). Within the MSA brain, high magnification microscopy revealed isSID signal within and adjacent to microglia (Fig. 3D, E, F). Additionally, microglia were observed in close contact with apparent isSID positive, IBA1 negative, cells (Fig 3E, Top panels). IBA1-positive sheet-like processes were intermingled within scattered punctate isSID labeling.

**Figure 3.**
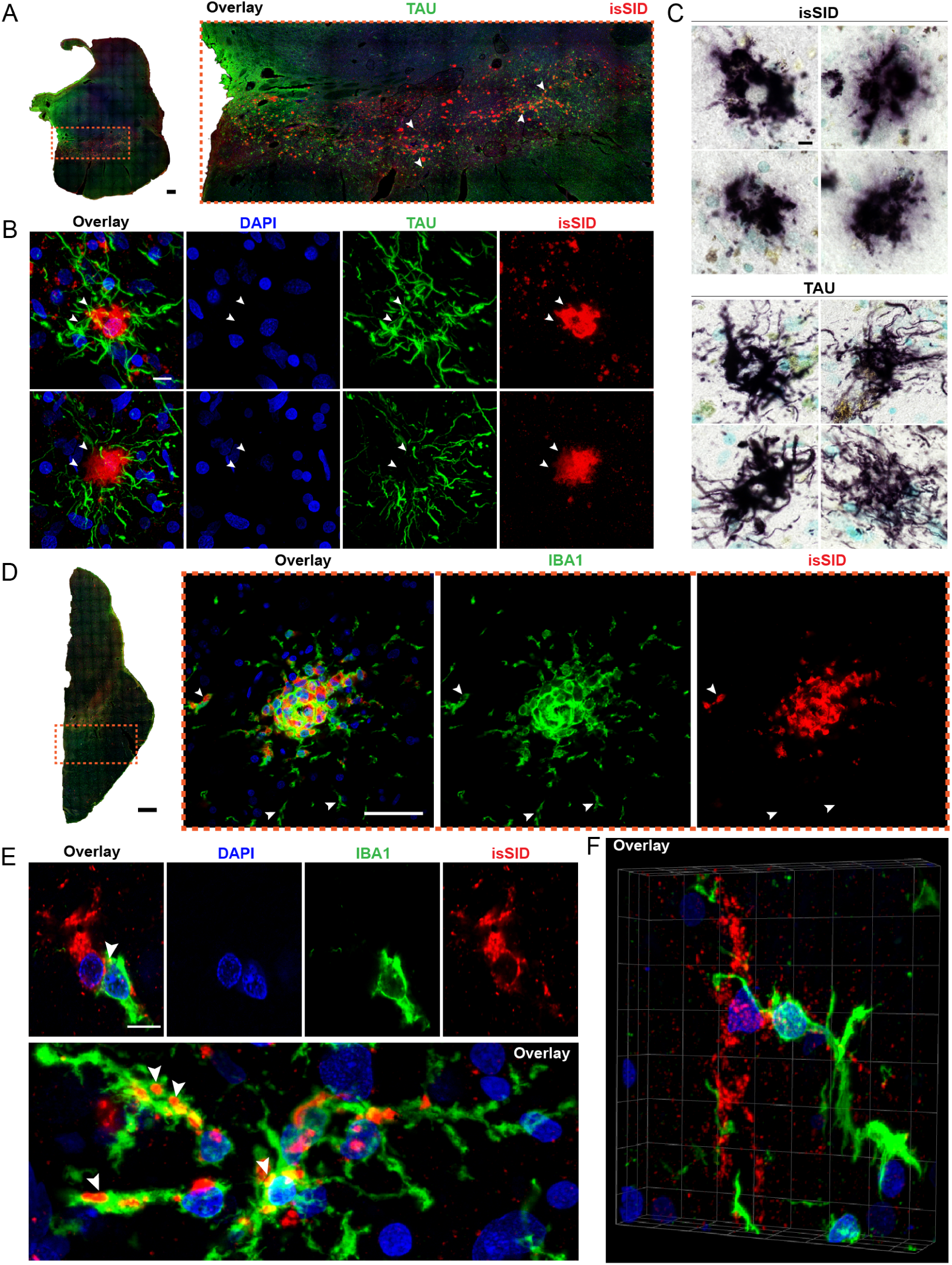
αsyn seeding activity is associated with tau pathology in PSP and microglia in MSA. **(A)** 10X overview of non-synucleinopathy control section, with an enlarged view of SN immunolabeled for Tau (green), isSID (red), and DAPI (blue, nuclei). Scale bar = 1 mm. **(B)** High-magnification images of tufted astrocytes showing a central pocket of seeding signal within the tau-positive pathological structure. Scale bar = 10 µm. **(C)** Comparison of isSID- and tau-DAB-processed sections from the SN showing similar morphological features of tufted astrocytes. Scale bar = 10 µm. **(D)** 10X overview of an MSA tissue section and high-magnification images of the cerebral peduncle (CP) immunolabeled for IBA1 (green), isSID (red), and DAPI (blue, nuclei). A cluster of microglia containing isSID positive species was observed, with IBA1-positive, isSID-negative microglia positioned along the periphery of the seed-positive region. Scale bar = 5 mm and 50 µm. **(E)** IBA1-positive microglial cells closely associated with seed-competent species were observed throughout the tissue, with overlapping signal observed at the site of contact (top panel), as well as clusters of microglia displaying discrete isSID-positive species localized along their cellular processes (bottom panel). Scale bar = 10 µm. **(F)** Three-dimensional volume reconstruction of an IBA-1 positive microglia cell in close proximity to an isSID-positive structure, demonstrating an extended microglial process oriented toward the seed-positive structure.

After characterizing isSID in synucleinopathy brain we then applied isSID-pSer129 multiplex labeling to striatal tissue sections from PD patients with mesencephalic neurons grafted into the putamen 18-month, 4, 16, and 27 years prior to death. At low magnification, strong pSer129 labeling was observed outside of the graft (Fig. 4A, 16-year graft shown, dotted line marks approximate graft location), but isSID was mostly ubiquitous diffuse labeling, with some overlap in the striatum and adjacent cortical tissue. Higher magnification within the graft revealed numerous pSer129-positive cells in the 16 year and 27-year grafts as previously described[45, 49, 50, 56]. Within pSer129-positive graft cells, strong isSID labeling was observed in Lewy body-like structures (Fig. 4B–D), as well as in diffuse and granular cytoplasmic patterns. We also observed isSID-negative Lewy bodies within the graft (Fig.4D, bottom-panels), but in those instances isSID signal was always identified proximally to the Lewy body. Lewy bodies and Lewy neurites within the host tissue (i.e., within the GP) were isSID positive, and had similar morphology to grafts (Fig. 4D, top-panel).

**Figure 4.**
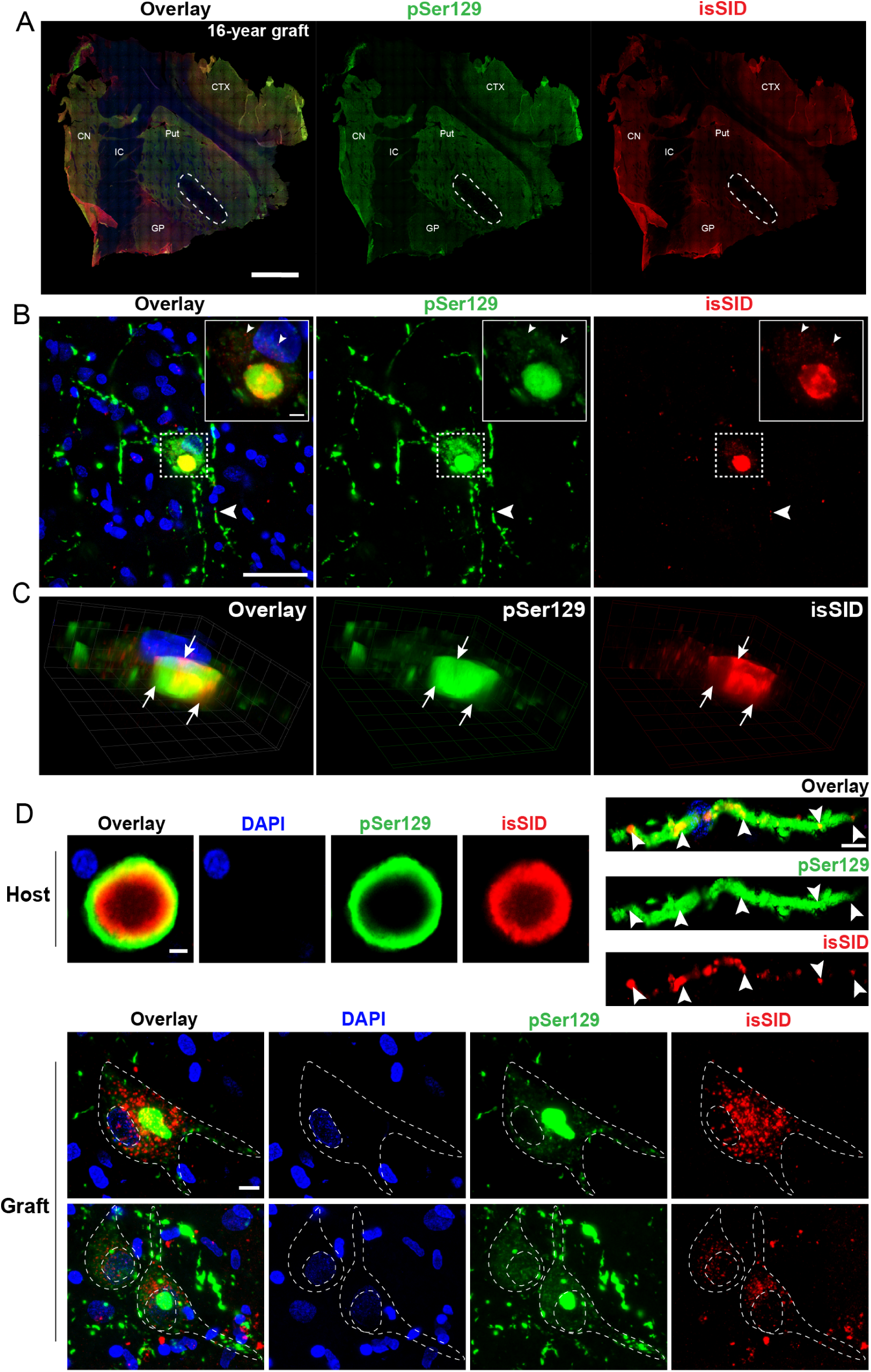
Seed-competent αsyn closely associates with pSer129 neuropathology in neuronal grafts years after transplantation. **(A)** 10X overview of 16-year putamen graft section immunolabeled for pSer129 (green), isSID (red), and DAPI (blue, nuclei). Scale bar = 5mm. (CN =Caudate Nucleus, IC = Internal Capsule, GP= Globus Pallidus, Put = Putamen, and CTX = Cortex.) **(B)** A prominent inclusion positive for both pSer129 and isSID surrounded by a pSer129 cloud with diffusely distributed isSID signal inside. Seed-competent αsyn was also detected within axonal processes extending from the grafted neuron. Scale bars = 5 µm and 50 µm. **(C)** Three-dimensional cross-section of the inclusion shown in panel B demonstrating localization of isSID signal within the pSer129-positive inclusion. The strongest isSID labeling corresponded to regions of reduced pSer129 immunoreactivity, whereas areas of intense pSer129 labeling exhibited comparatively weaker isSID signal. **(D)** Comparison of host- and graft-derived αsyn pathology. The host-derived αsyn pathology, including a Lewy body and Lewy neurite, exhibited characteristic pSer129-positive morphology, with isSID signal localized within or along the pSer129-positive structures. In contrast, graft-derived neurons display diffuse, whole-cell pSer129 immunoreactivity. Within these grafted neurons, isSID-positive seed signal appears as discrete puncta associated with and surrounding pSer129-positive inclusions. Scale bars = 5 µm and 10 µm.

Some, but not all, grafted cells contain neuromelanin and express TH. To determine if seeding was occurring within TH expressing grafted cells, we multiplex labeled isSID and TH. Results show abundant TH at the site of implantation, with diffuse isSID, and little overlap at low magnification (Fig. 5A). However, closer magnification identified strong TH labeling in the grafts, with many TH/isSID positive cells apparent (Fig. 5B). TH-positive cells were isSID positive, showing diverse labeling morphologies, including dense Lewy-body-like labeling, and diffuse/granular (Fig. 5B, C, D). We also observed many cells that lacked isSID labeling (Fig, 5D). Notably, some, but not all isSID signals were associated with neuromelanin within the grafts (Fig. S3).

**Figure 5.**
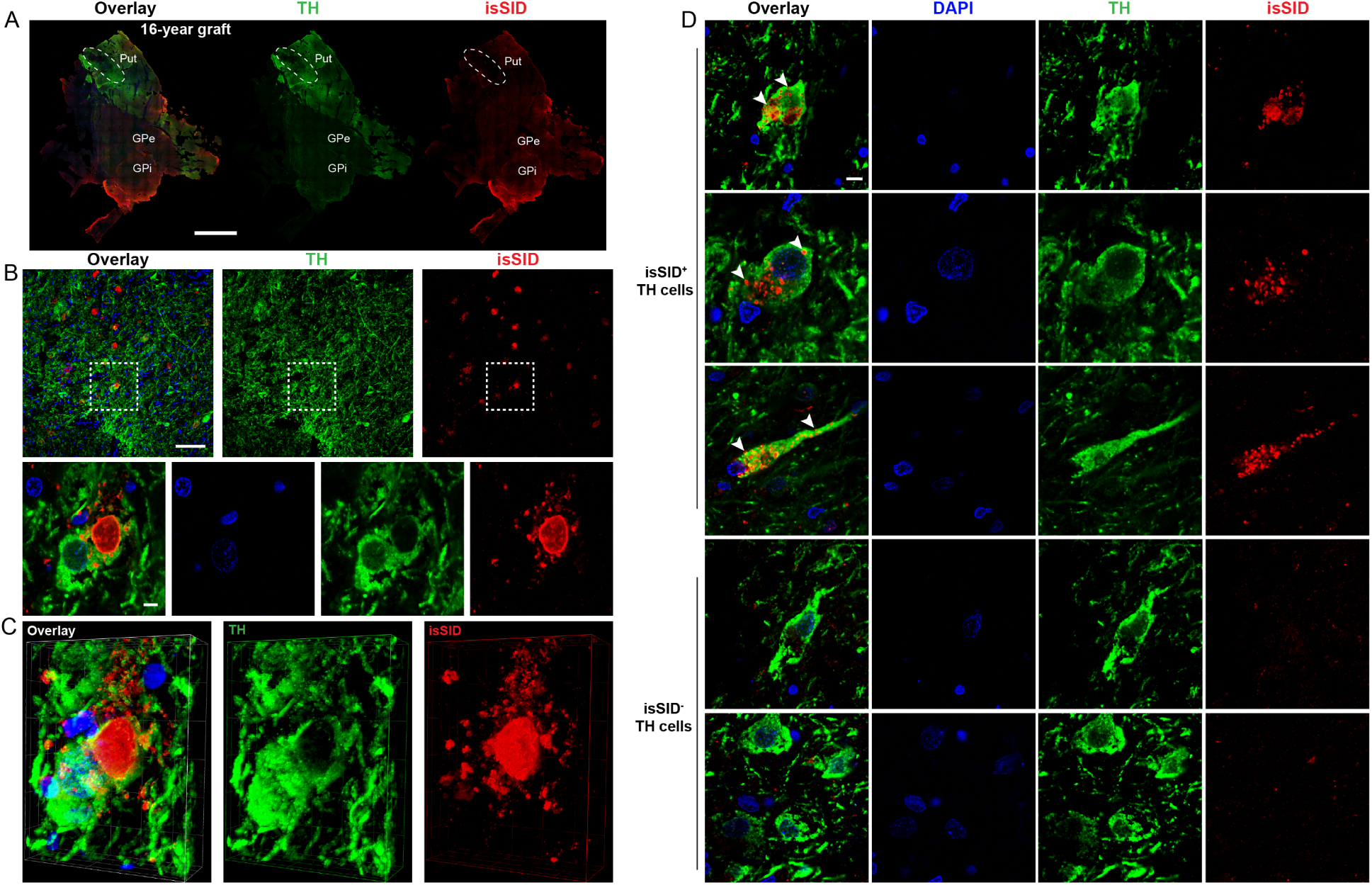
Seed-competent αsyn is detectable in grafted dopamine neurons. **(A)** 10X overview of 16-year putamen graft section immunolabeled for TH (green), isSID (red), and DAPI (blue, nuclei). Scale bar = 5 mm. (GPe= External Segment of Globus Pallidus, GPi = Internal Segment of Globus Pallidus, and Put = Putamen). **(B)** Low-magnification of the grafted area illustrating the heterogenous distribution of seed-competent αsyn among TH-positive cells, with seed-positive profiles not uniformly associated with TH signal. A representative TH-positive cell containing isSID signal is shown at higher magnification, with the seed signal concentrated in a perinuclear, inclusion-like pattern reminiscent of a Lewy body and encircled by TH immunoreactivity. A cluster of smaller punctate signals can be observed surrounding the seed-positive signal. Scale bar = 10 µm and 100 µm. **(C)** Three-dimensional volume reconstruction of the representative cell shown in panel B highlighting the deep TH-positive pocket surrounding the isSID inclusion. **(D)** Representative panel of isSID-positive and - negative TH cells. Positive seed signal was observed in discrete, localized clusters distributed throughout the entirety of the TH-positive cell. Scale bar = 10 µm.

To establish the precise origin of seeding within the grafts, we then multiplex labeled graft specimens for TH, isSID, and pSer129 (Fig 6, 27-year graft shown, except Fig 6C. bottom two panels). This approach allowed us to directly confirm whether grafted dopamine neurons contained isSID positive Lewy bodies. Results show overall distribution of TH, pSer129, and isSID as before (Figure 6A). Higher magnification of the grafts (16 and 27-year) revealed numerous examples of grafted neurons that were both isSID and pSer129 positive (Fig. 6B-C). Concentric ring like Lewy bodies were often isSID positive, however some grafted cells contained both pSer129 and isSID, with minimal direct overlap (Figure 6C, bottom two panels). We did not observe any pSer129-postive grafted cells that lacked isSID.

**Figure 6.**
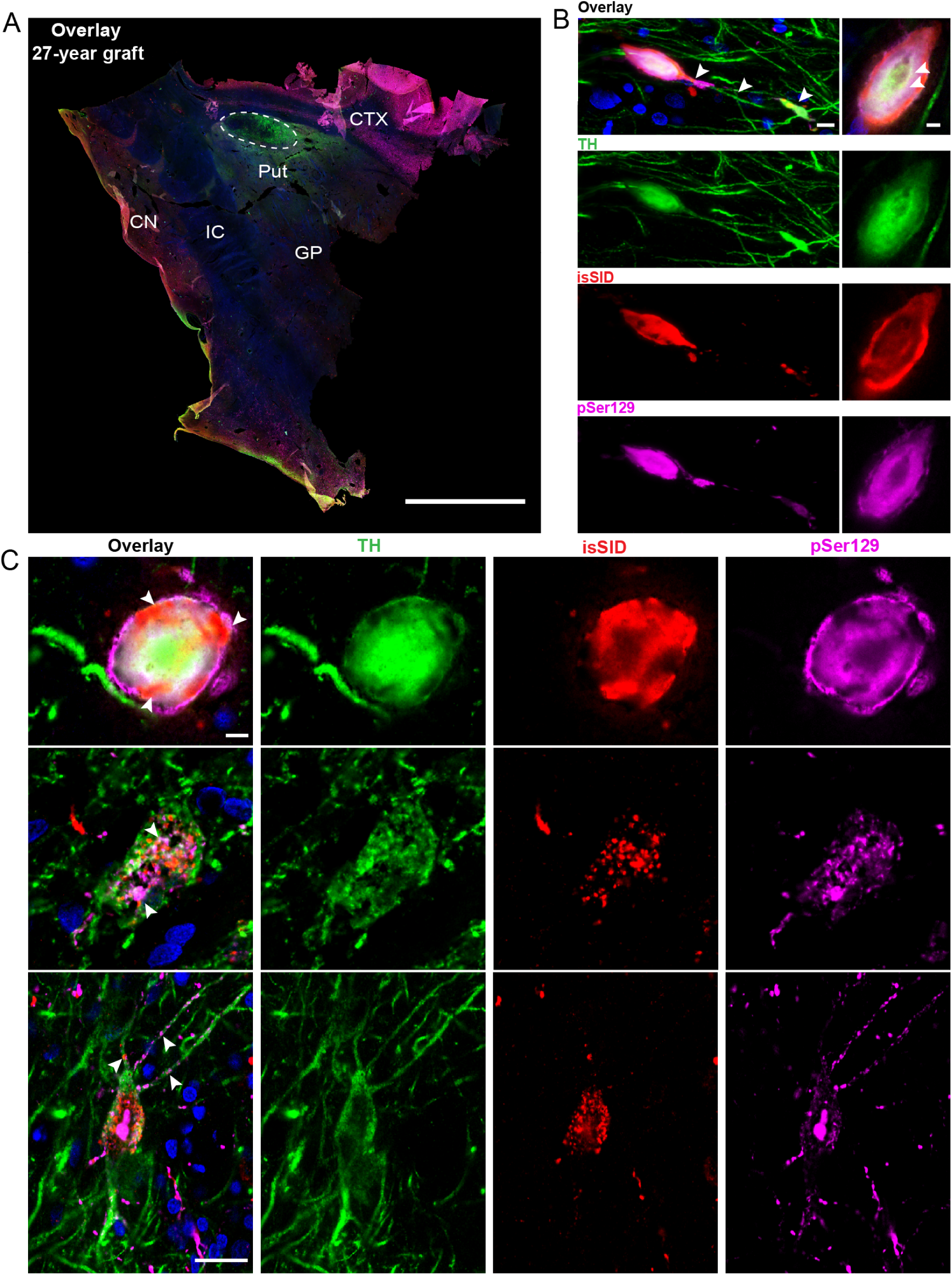
Seeding occurs in grafted dopamine neurons bearing LP. **(A)** 4X overview of 27-year putamen graft section immunolabeled for TH (green), isSID (red), pSer129 (pink), and DAPI (blue, nuclei). Scale bar = 10 mm. CN =Caudate Nucleus, IC = Internal Capsule, GP= Globus Pallidus, Put = Putamen, and CTX = Cortex. **(B)** A single TH-positive grafted neuron exhibiting overlapping isSID and pSer129 signal. Higher-magnification imaging revealed an inclusion-like accumulation of pSer129 and isSID signal characterized by a distinct central clearing within the pSer129-postive structure, reminiscent of Lewy body-like pathology. Scale bar = 10 µm and 20 µm. **(C)** Panel of triple labeling in the 27-year (top panel) and 16-year (bottom two panels) putamen grafts. A more mature pSer129-positive inclusion is observed in the 27-year graft compared with the 16-year graft, with a noticeable absence of smaller punctate isSID-positive structures surrounding the inclusion. Scale bar = 10 µm and 50 µm.

Younger grafts, 18 month and 4 years, lacked pSer129, but were isSID positive, particularly the few neuromelanin containing cells and amorphous isSID signal outside of TH-cells (Fig. 7A and B, respectively). isSID in 18 month grafted cells, was granular, and lacked the compact concentric ring morphology of Lewy bodies (Fig. 7A, Bottom Panels). pSer129 and isSID strongly labeled outside grafted areas, both within the putamen, caudate, globus pallidus, and cortex (Fig. 7).

**Figure 7.**
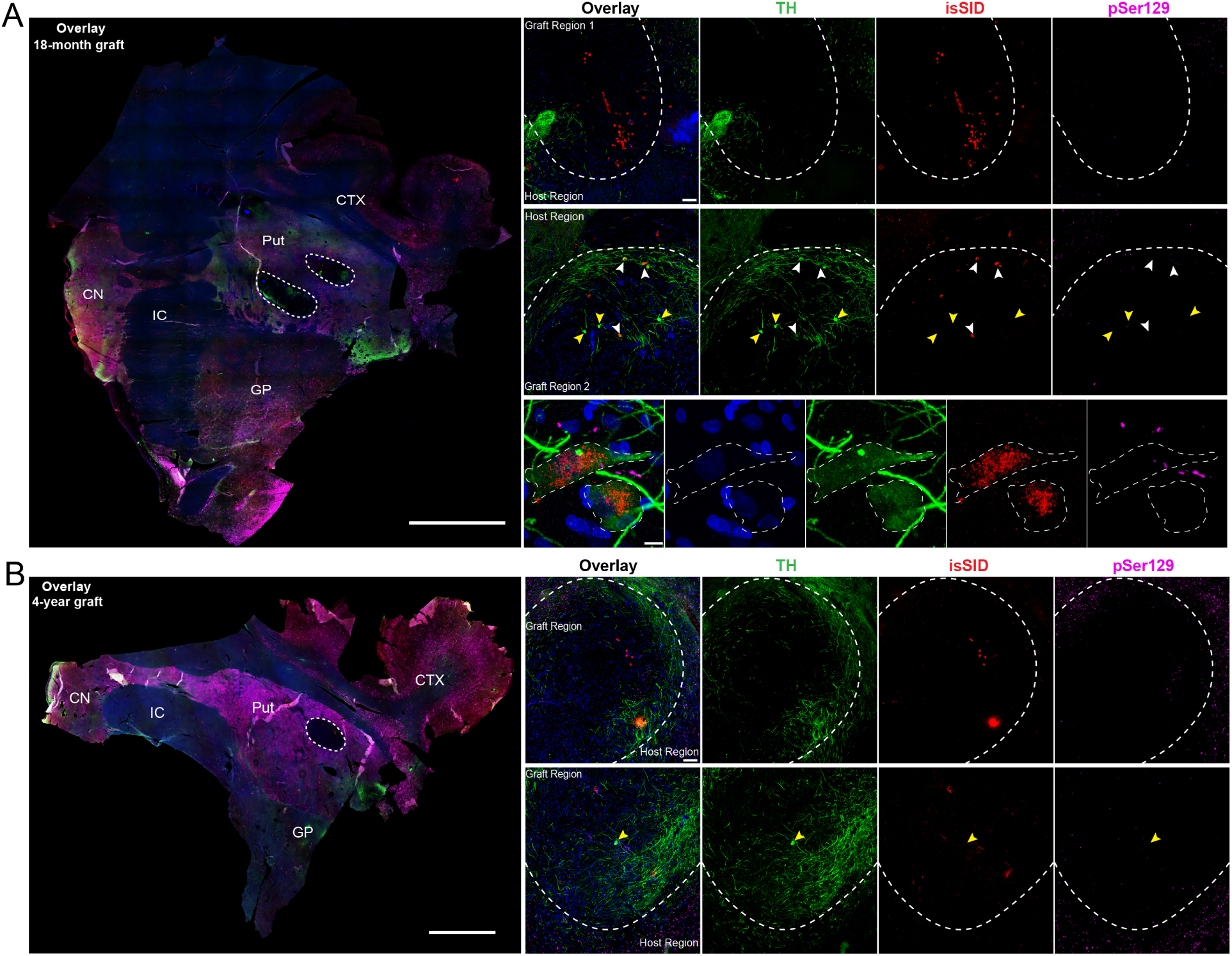
Younger grafts display amorphous isSID signal with minimal TH overlap. **(A)** 4X overview of 18-month putamen graft section immunolabeled for TH (green), isSID (red), pSer129 (pink), and DAPI (blue, nuclei). Scale bar = 10 mm. (CN =Caudate Nucleus, IC = Internal Capsule, GP= Globus Pallidus, Put = Putamen, and CTX = Cortex.) Top grafted site displaying isSID signal independent of TH-positive structures (top panel). Bottom grafted site highlighting heterogenous environment of grafted region, with both isSID-positive (white arrows) and -negative (yellow arrows) TH cells (middle panel). High-magnification image of two isSID-positive TH cells displaying granular clusters of isSID signal that lack detectable pSer129 signal (bottom panel). Scale bar = 10 µm and 100 µm. **(B)** 4X overview of 4-year graft section immunolabeled for TH (green), isSID (red), pSer129 (pink), and DAPI (blue, nuclei). Scale bar = 5 mm. (CN =Caudate Nucleus, IC = Internal Capsule, GP= Globus Pallidus, Put = Putamen, and CTX = Cortex.) Area one of the grafted site displaying a similar independent isSID signal pattern as observed in the top grafted site of 18-month (top panel). Area two of the grafted site with a single TH cell lacking detectable isSID and pSer129 signal (bottom panel). Scale bar = 100 µm.

## Discussion

The observation that LP can develop over time in grafted immature neurons (typically starting a decade after transplantation surgery) represents evidence that αsyn spreads in the human PD brain. However, other mechanisms, such as inflammation, may also be involved. Here, we show that active seeding occurs prior to and alongside LP formation. Indeed, we saw robust seeding in grafts that largely overlapped with LP markers (e.g., pSer129), and in agreement with previous reports[31, 42, 48, 49] not all grafted cells contained seeding and LP. Spontaneous, idiopathic αsyn seeds have only been documented in human brain. Graft neurons are young, without known PD genetic risk background, and therefore seeding inside them is unlikely to be a *de novo* idiopathic event. Although these findings cannot fully exclude alternative mechanisms, they, together with preclinical studies demonstrating the transfer of host-derived αsyn to grafted fetal nigral neurons following transplantation into transgenic αsyn-overexpressing mice[33] or following AAV-mediated αsyn expression in rats[43], provide strong support for a mechanism in which pathological αsyn seeds originating in host neurons promote the propagation of disease-associated pathology into grafted neurons.

Seeding was robust in 16-year-old and 27-year-old grafts, but less so in younger grafts, consistent with progressive LP development in the grafts[42, 48]. Although young grafts (18 month and 4 year) lacked overt pSer129-positive LP, we did find isSID-positive granular structures within the few neuromelanin-containing cells of the 18 month graft, as well as large amorphous structures within grafted area, but not overlapping with the TH-immunopositive neurons. This suggests that seeding in these cases can occur but more time is required for sufficient levels of LP to be visualized. This contrasts with older grafts, where isSID appeared as large Lewy body-like ring inclusions, and discrete punctate inclusions found both in perinuclear space and in apparent Lewy neurites. isSID in the host tissue was morphologically similar to structures in the grafts.

Our multiplex labeling of pSer129 and isSID confirmed seeding within LP in the PD/DLB brain and within GCIs in the MSA brain. Interestingly, isSID partially overlapped with LP and GCIs suggesting that only a subpopulation of αsyn pathology was actively seeding. Alternatively, the post-translational pSer129 modification may inhibit αsyn aggregation[27] which may explain incomplete isSID overlap with pSer129-positive LP, as pSer129 might inhibit αsyn binding *in situ*. In future studies additional αsyn antibodies could help further define actively seeding αsyn species, or the location of active nucleation centers in the brain. Indeed, isSID and pSer129 showed a similar spatial segregation in PFF models with seeding occurring discretely along pSer129 αsyn aggregates[10].

We also observed isSID αsyn binding outside the context of αsyn pathology (i.e., not overlapping with pSer129-positive inclusions). For example, isSID labeled tufted astrocytes in the PSP midbrain [57], consistent with previous observations of isSID reactive tau-tangles[57] and known interactions between αsyn and tau[17, 28, 30]. αSyn also bound to neuromelanin in midbrain and in the grafts, regardless of diagnosis and synucleinopathy status. Previous reports found that isSID labeling in a non-synucleinopathy brain[57], which the authors concluded could represent incidental pathology, but our observations suggest this labeling may be driven by αsyn affinity for neuromelanin. Indeed, they found isSID reactivity in melanated cells corresponding to total αsyn staining (i.e., clone 42)[57], suggesting isSID labeled neuromelanin-containing endogenous αsyn. Interestingly, αsyn sequesters into neuromelanin granules before LP formation[21], and neuromelanin accumulation in animal models can result in concentric Lewy-body like inclusions in midbrain neurons [8, 9, 21, 32]. Our observation that αsyn specifically binds neuromelanin in the human brain supports the hypothesis that neuromelanin acts as a catalyst/nucleation center for αsyn aggregates[22, 32, 64]. Neuromelanin organelles are autolysosome-like granules storing proteins, and oxidized lipids covalently attached to a metallic core, and isSID labeling may represent an affinity of αsyn for stored lipids[32]. Broadly, αsyn interaction with lipids is increasingly viewed as central to the pathogenesis of synucleinopathy, although many details remain unclear [20, 41]. In addition to neuromelanin labeling, we found amorphous isSID accumulations in PD, MSA, PSP, and graft specimens. For many of these, the molecular makeup of the structures is unclear. However, we were able to establish that those strong isSID reactivity in MSA brain were microglial nodules, which occur in degenerating white matter tracts of MSA brain, consistent with local microgliosis acting to remove αsyn seeds[36, 38].

There are some limitations for our studies. Although we demonstrate active seeding occurred in the grafted neurons, inflammation or other host factors still might play a role in the development of LP in the grafted neurons. Furthermore, post-mortem specimens cannot prove directionality or the kinetics of seed transfer. All human brain specimens had confirmed movement disorders, and were elderly patients, limiting our conclusions about the role of αsyn-neuromelanin binding, and its possible disease relevance. Future studies should assess brain specimens from neurologically intact, young, patients, to determine whether αsyn affinity for neuromelanin is innate, or disease relevant. Lastly, we validated isSID in human synucleinopathy brain and then applied to grafts; follow-up studies on some interesting observations (e.g., seeding in microglial nodules) in the synucleinopathy brain are warranted.

In conclusion, αsyn seeding precedes LP formation, in young, healthy neurons grafted to the PD brain, with early seeding being spatially associated with neuromelanin granules. While post-mortem tissue do not establish direction or kinetics, the present findings strengthen the case that αsyn aggregates propagate in the human PD brain and that preventing spread can be relevant both to the development of therapies that slow disease progression and to promoting the long-term efficacy of cell-replacement therapies.

