## Supplemental Information for "α-Synuclein Seeding in Dopaminergic Neurons Transplanted into the Putamen of Parkinson’s Disease Patients"

Table of Contents:

Figure S1. Human and mouse αsyn bind to Lewy pathology in the PD brain

Figure S2. αSyn exhibits preferential labeling of some neuromelanin associated structures.

Figure S3. αsyn binds to a select subset of neuromelanin-associated structures in grafted neurons

Figure S4. αsyn seeding activity is not associated with beta-amyloid (Aβ) pathology in PSP or astrocytes in MSA.

Figure S5. Mouse-on-Mouse (M.O.M) blocking does not effectively prevent interference from mouse-derived secondary antibody for isSID.

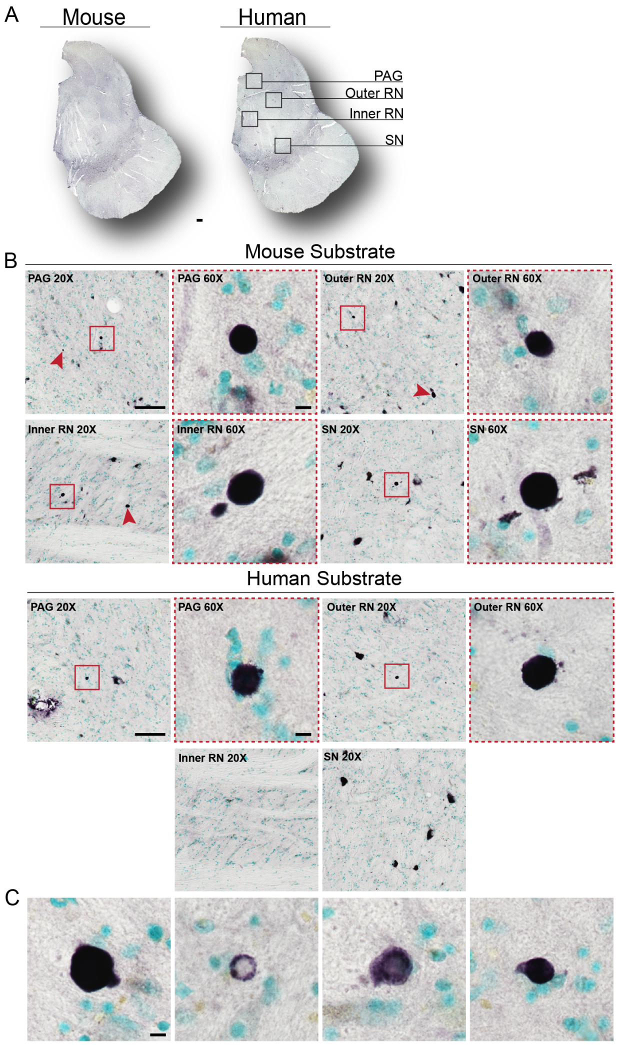


Figure S1. Human and mouse αsyn bind to Lewy pathology in the PD brain (A) 10X overview of matched midbrain DAB-processed tissue sections using both mouse and human substrate. Scale bar = 1 mm. (B) High- and low- magnification of periaqueductal grey (PAG), outer red nucleus (outer RN), inner red nucleus (inner RN), and substantia nigra (SN) highlighting regional abundance of seeding detection between both substrates. Scale bar = 10 μm and 100 μm. Mouse substrate detected more seed-competent species per region of interest (ROI) as indicated by the red arrows. (C) Panel of seed-competent species detected with human substrate in other areas of the midbrain. Some show complete dark labeling, whereas others are fainter and only show signal detection in certain areas of the structure. Scale bar = 10 μm.

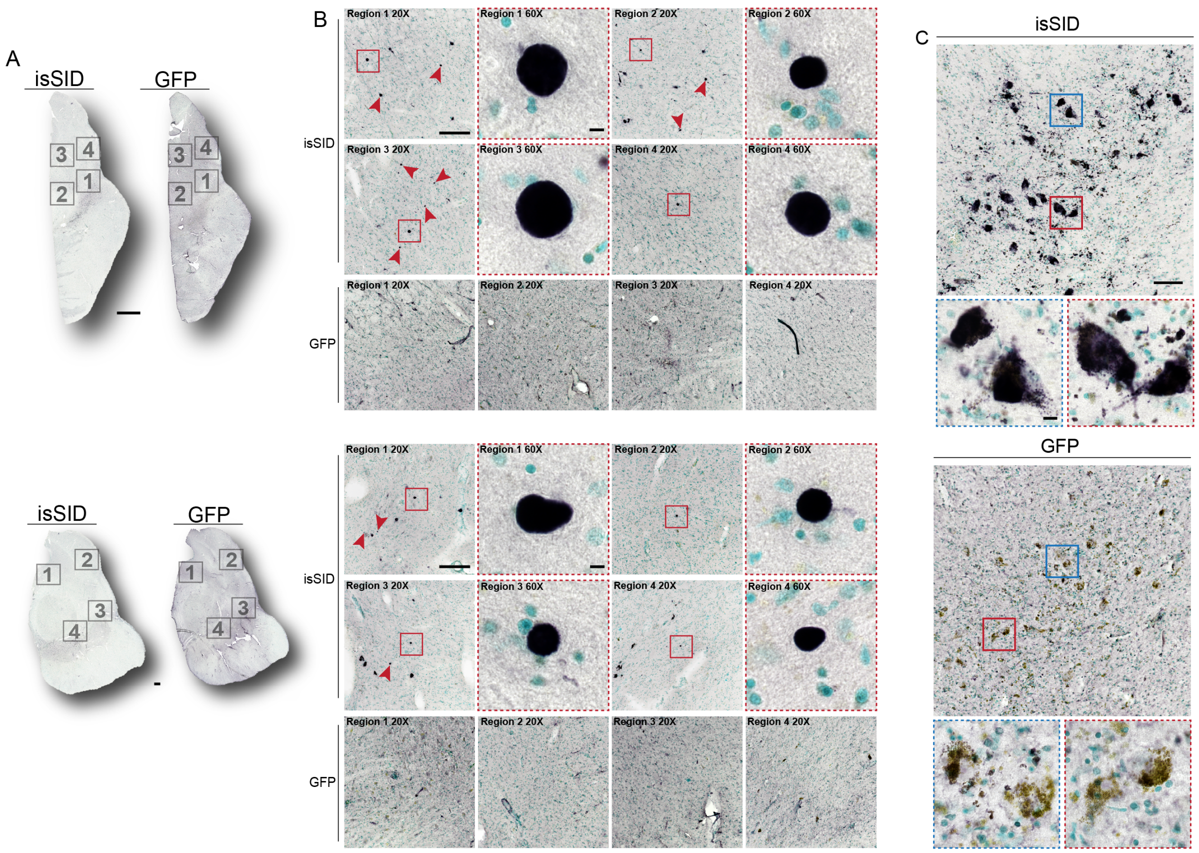


Figure S2. αSyn exhibits preferential labeling of some neuromelanin associated structures. (A) 10X overview of two specimens processed with isSID and GFP simultaneously. Scale bar = 1 mm and 5 mm. (B) High- and low-magnification of individual regions (1-4) of both isSID and GFP for the two specimens. Seed-competent structures are only detectable in isSID condition and not in the GFP condition. Red arrows indicate additional isSID positive structures that were identified in that region of interest. Scale bar = 10 μm and 100 μm. (C) Neuromelanin staining patterns in the substantia nigra under both conditions. Selected regions were imaged at higher magnification as indicated by the red and blue boxes. Pigments labeled with isSID appear noticeably darker and are accompanied by the presence of smaller punctate structures inside and surrounding the larger labeled pigment. Scale bar = 10 μm and 100 μm.

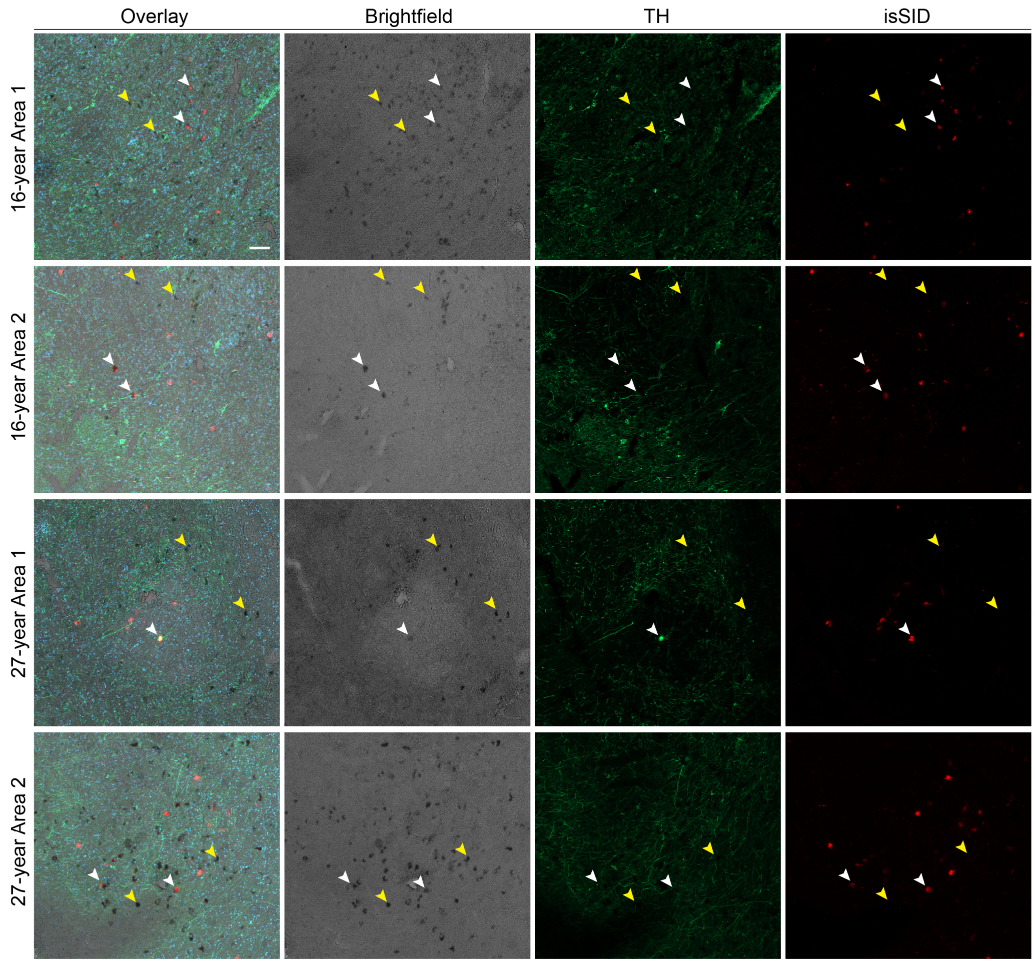


Figure S3. αsyn binds to a select subset of neuromelanin-associated structures in grafted neurons. Overlay of brightfield and fluorescent TH and isSID images demonstrate the distribution of neuromelanin-associated structures and seed signals. isSID signal was observed in a subset of pigmented cells across both 16 and 27-year grafts (white arrows), whereas many neuromelanin cells lacked detectable isSID signal (yellow arrows). Scale bar = 100 µm.

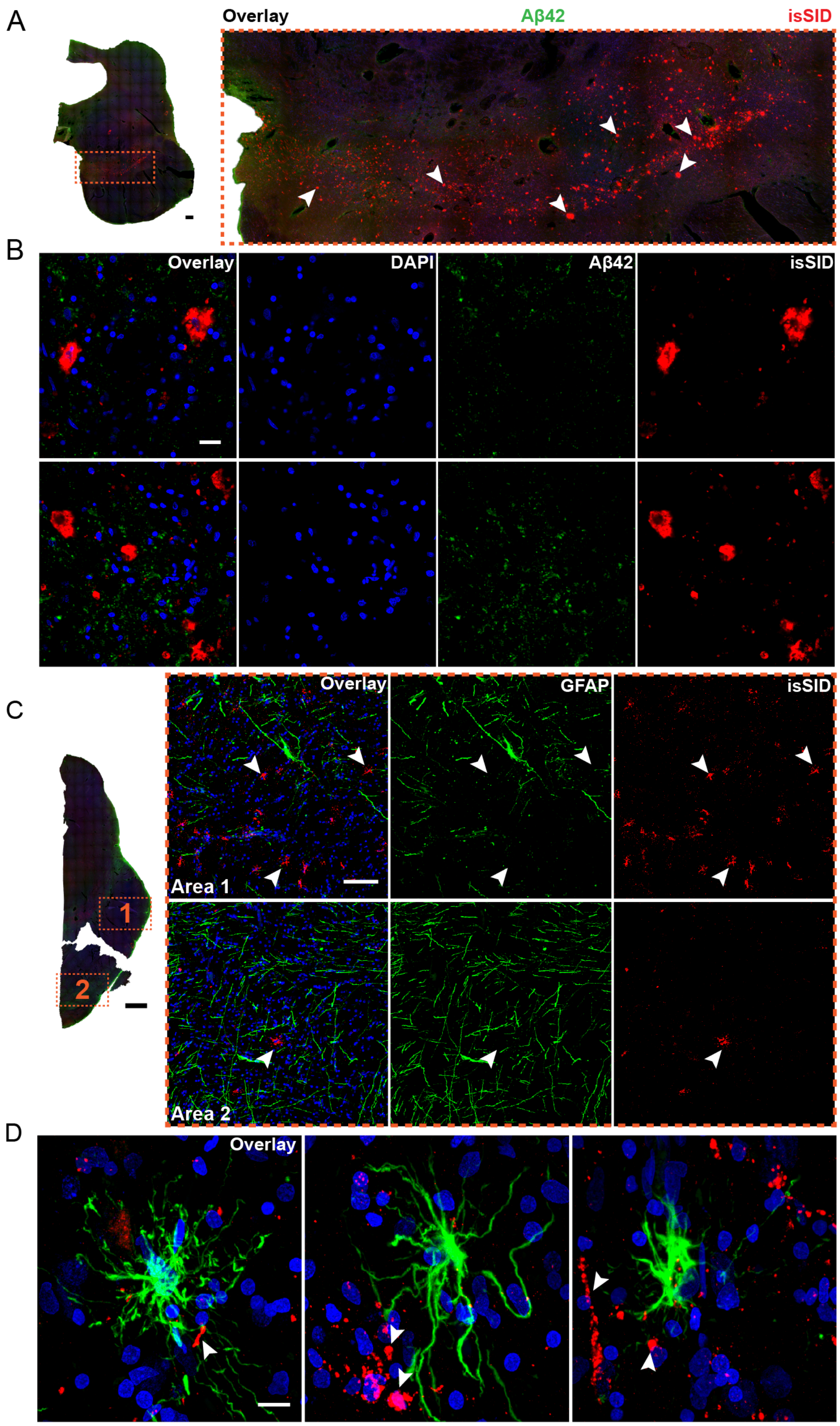


Figure S4. αsyn seeding activity is not associated with beta-amyloid (Aβ) pathology in PSP or astrocytes in MSA. (A) 10X overview of non-synucleinopathy control section, with an enlarged view of SN immunolabeled for Aβ42 (Abcam Cat# ab201060, RRID:

AB_2818982 (green)), isSID (red), and DAPI (blue, nuclei). Scale bar = 1 mm. (B) High-magnification images of the SN region revealed minimal Aβ42 signal, which appeared primarily as diffuse background staining without apparent discrete plaque pathology. Scale bar = 20 μm. (C) 10X overview of MSA case immunolabeled for GFAP (Abcam "EPR1034Y" Cat# ab68428, RRID: AB_1209224(green)), isSID (red), and DAPI (blue, nuclei). Scale bar = 5 mm. Low-magnification images of cerebral peduncle areas one and two highlight a spatially inverse distribution of GFAP and isSID signal. Areas of increased isSID signal exhibited lower GFAP signal, while regions with higher GFAP signal showed relatively less isSID signal. Scale bar = 100 μm. (D) Representative high-magnification images of whole astrocytes revealed spatial separation of GFAP and isSID signal, with the most prominent isSID-positive structures not overlapping with GFAP-positive astrocytes. Scale bar = 10 μm.

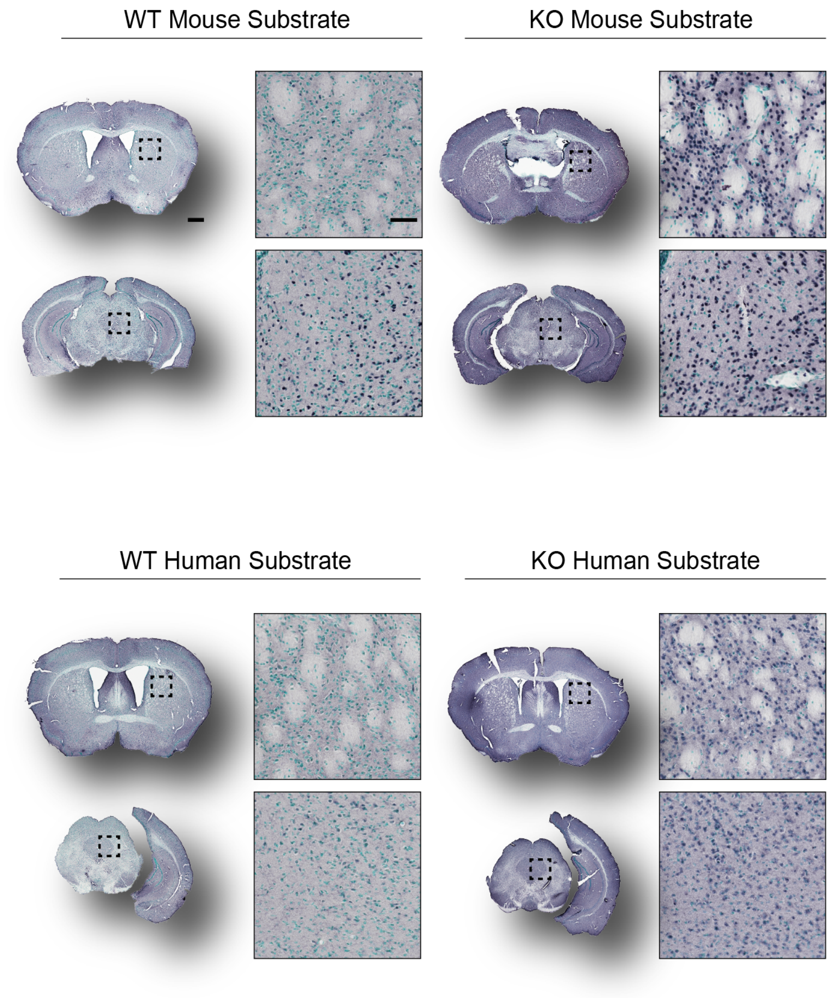


Figure S5. Mouse-on-Mouse (M.O.M) blocking does not effectively prevent interference from mouse-derived secondary antibody for isSID. 10X overview of wild-type (WT) and αsyn knockout (KO) mouse striatum and midbrain sections processed with mouse and human substrate following pretreatment with M.O.M blocking reagent (Vector Laboratories Cat# MKB-2213, RRID: AB_2336587). Higher magnification images of the striatum and the periaqueductal grey (PAG) region of the midbrain revealed strong, ubiquitous background labeling in both WT and KO sections across conditions, which was partially eliminated by using a goat-derived secondary (Fig 1E). Scale bar = 50 μm and 1 mm.
